# Transcriptome analysis reveals that fertilization with cryopreserved sperm downregulates genes relevant for early embryo development in the horse

**DOI:** 10.1101/558304

**Authors:** JM Ortiz-Rodriguez, C Ortega Ferrusola, MC Gil, FE Martín Cano, G Gaitskell-Phillips, H Rodríguez-Martínez, K Hinrichs, A Álvarez-Barrientos, A Román, FJ Peña

**Affiliations:** Department of Veterinary Physiology and Pharmacology, College of Veterinary Medicine & Biomedical Sciences, Texas A&M University, College Station, Texas.; Department of Biochemistry and Molecular Biology, University of Extremadura, Badajoz, Spain; STAB, University of Extremadura, Badajoz, Spain; Laboratory of Equine Reproduction and Equine Spermatology, Veterinary Teaching Hospital, University of Extremadura, Cáceres, Spain

**Keywords:** equine, sperm, cryopreservation, embryo, RNAseq, transcriptome

## Abstract

Artificial insemination with cryopreserved sperm is a major assisted reproductive technology in many species. In horses, as in humans, insemination with cryopreserved sperm is associated with lower pregnancy rates than those for fresh sperm, however, direct effects of sperm cryopreservation on the development of resulting embryos are largely unexplored. The aim of this study was to investigate differences in gene expression between embryos resulting from fertilization with fresh or cryopreserved sperm. Embryos were obtained at 8, 10 or 12 days after ovulation from mares inseminated post-ovulation on successive cycles with either fresh sperm or frozen-thawed sperm from the same stallion, providing matched embryo pairs at each day. RNA was isolated from two matched pairs (4 embryos) for each day, and cDNA libraries were built and sequenced. Significant differences in transcripts per kilobase million (TPM) were determined using (i) genes for which the expression difference between treatments was higher than 99% of that in the random case (P < 0.01), and (ii) genes for which the fold change was ≥ 2, to avoid expression bias in selection of the candidate genes. Molecular pathways were explored using the DAVID webserver, followed by network analyses using STRING, with a threshold of 0.700 for positive interactions. The transcriptional profile of embryos obtained with frozen-thawed sperm differed significantly from that for embryos derived from fresh sperm on all days, showing significant down-regulation of genes involved in biological pathways related to oxidative phosphorylation, DNA binding, DNA replication, and immune response. Many genes with reduced expression were orthologs of genes known to be embryonic lethal in mice. This study, for the first time, provides evidence of altered transcription in embryos resulting from fertilization with cryopreserved spermatozoa in any species. As sperm cryopreservation is commonly used in many species, including human, the effect of this intervention on expression of developmentally important genes in resulting embryos warrants attention.

## INTRODUCTION

Cryopreservation is a common procedure in assisted reproductive technology, in both humans and the animal breeding industry [1-3]. Cryopreserved sperm are routinely used for artificial insemination (AI), in vitro fertilization (IVF) and intracytoplasmic sperm injection (ICSI). However, it is clear that sperm cryopreservation methods are currently sub-optimal, as pregnancy rates with cryopreserved sperm are lower than those with fresh sperm in humans and horses [4], among other species. Cryopreservation leads to extensive damage of sperm cell membranes and causes metabolic and functional alteration of sperm [5, 6], particularly of their mitochondria [7-9]. Cryopreservation may alter sperm DNA [10]; recently, specific cryodamage to sperm genes and transcripts have been reported [11, 12], even in samples with good sperm motility post thaw and in the absence of detectable DNA fragmentation. The sperm DNA is epigenetically programmed to regulate embryonic gene expression, and changes to this epigenome cause developmental disregulation [13]. Cryopreservation has been found to significantly change the sperm DNA methylome, as well as to alter expression of epigentic-related genes such as methyltransferases (Aurich, Zang). Cryopreservation of sperm imposes oxidative stress and redox deregulation in spermatozoa, leading to the presence of toxic adduct-forming compounds such as 4- hydroxynonenal (4-HNE) in sperm membranes [14]. Moreover, mitochondria of spermatozoa surviving cryopreservation show increased production of reactive oxygen species [7, 8, 15]. Signaling pathways crucial to normal embryo development are sensitive to perturbations of endogenous redox state, and are also susceptible to modulation by reactive oxygen species [16]. Thus, fertilization by damaged spermatozoa may impact early embryo development and even have effects that appear later in the life of the offspring [17].

Moreover, appreciation of the contribution of sperm to embryo development has evolved from the concept that the only role of sperm at fertilization is to introduce the male genome into the egg. Sperm carry a myriad of small noncoding RNAs with potential roles in early embryo development [18, 19]. Notably, sperm carry the activating factor PLCζ, which triggers calcium oscillations that induce oocyte activation [20, 21], and alterations in frequency and amplitude of post-fertilization calcium oscillations can affect the phenotype of the resulting embryo into adulthood [22]. Thus, there are extensive pathways by which cryopreservation of sperm could alter the development of the fertilized ooctye and embryo.

Despite the widespread use of cryopreserved sperm, and the known decrease in pregnancy rates with its use, little direct information is available on the effect of sperm cryopreservation on development of the resulting embryo. Recent advances in transcriptome amplification and next generation sequencing provide the ability to obtain the full transcriptome of individual embryos [23], thus offering a basis for studies on differences in gene expression associated with fertilization with cryopreserved sperm. In the present study, we analyzed the transcriptome of equine embryos produced with fresh or frozen-thawed sperm, to determine the impact of sperm cryopreservation on gene expression during early equine embryo development.

## MATERIAL AND METHODS

### Animals and experimental design

Animals were maintained according to European laws and regulations, and all experimental procedures were reviewed and approved by the Ethical committee of the University of Extremadura, Cáceres, Spain. Six mares and two stallions of known fertility were used for this study. Each mare was assigned a day of embryo recovery (8, 10 or 12 days post ovulation) and on successive cycles was assigned to be inseminated with fresh or frozen-thawed sperm from the same stallion, to provide matched embryo pairs for that day of embryo development. The mares were treated with a prostaglandin analogue to shorten the luteal phase and were monitored daily by transrectal ultrasonography. When a follicle of at least 35 mm diameter was detected in the absence of luteal tissue, with marked uterine edema and low cervical tone, mares received 2,500 IU of hCG i.v.. The follicle was monitored by transrectal ultrasonography every 6 h thereafter to detect the time of ovulation. Mares were inseminated immediately once ovulation was detected, with a minimum of 100 million either fresh sperm or frozen-thawed sperm, from the same stallion. Embryos were obtained by uterine lavage on the designated day after ovulation. For each embryo day, two embryos produced with fresh sperm, designated FRSH embryos, and 2 embryos produced with frozen-thawed sperm, designated CRYO embryos were obtained. Embryos were snap-frozen in liquid N_2_ and stored at −80°C until analysis. Previous clinical reports indicated that there is no a significant effect in the rate of embryonic vesicle growth between mares inseminated with fresh or frozen-thawed sperm if both are inseminated post-ovulation [24].

### Isolation of RNA

Total RNA was isolated from the embryos using the PicoPure™ RNA Isolation Kit (Catalog number KIT0204, Thermofisher) following the manufacturer’s instructions. RNA concentration and quality were assessed by automatic electrophoresis using 2100 Bioanalyzer (Agilent Technologies, Santa Clara, CA, USA).

### RNA-seq analysis

cDNA libraries were built using an IonTorrent S5/XL sequencer (Thermo Fisher Scientific, Waltham, MA USA). The raw reads were aligned to a horse transcriptome generated using ENSEMBL (Equ Cab 2 version) in the Torrent server with proprietary ThermoFisher algorithms. Then, custom scripts were used to transform reads into transcript counts, and transcripts per kilobase million (TPM) scores for each gene were retrieved. A gene was considered expressed if the reads per kilobase or transcript model per million mapped reads was > 0.4. In order to evaluate gene expression differences between treatments (FRSH or CRYO embryos), we calculated two thresholds: first, we calculated the random TPM differences between FRSH and CRYO embryos by permutation of the TPM gene scores. Then we chose the genes whose expression difference between the two conditions was higher than in 95% (P<0.05) or in 99% (P<0.01) of the random cases. As a second score, we used a fold change ≥ 2 as a threshold in order to avoid expression biases in the selection of the candidate genes.

### Gene Ontology and pathway analysis

The annotations of the candidate genes selected after the RNA-seq analyses were explored to detect significant differences in molecular pathways between treatments. Specifically, the DAVID webserver [25] was used to retrieve the terms (gene ontology, up-expressed tissues, KEGG and reactome pathways, protein-protein interactions, etc) with significant over-presence of the candidate genes, using a false discovery rate (FDR) < 0.05. We used the human genome as reference for the analysis because of its increased depth in terms of annotation.

### Network analysis

STRING [26] was used to analyze the internal structure of the functional network obtained using the candidate genes. Data included co-expression, genetic fusion, co-occurrence or protein-protein interactions, among others. A high threshold (0.700) was selected for positive interaction between a pair of genes.

## RESULTS

A total of 12 conceptuses were analysed (2 FRSH and 2 CRYO at each day). An average of 29,196 transcripts per embryo were obtained.

### Day-8 embryos

In Day-8 CRYO embryos, 100 transcripts showed increased abundance and 157 transcripts showed decreased abundance in respect to FRSH embryos of the same age from the same stallion and mare (Fig. 1).

**Figure 1.**
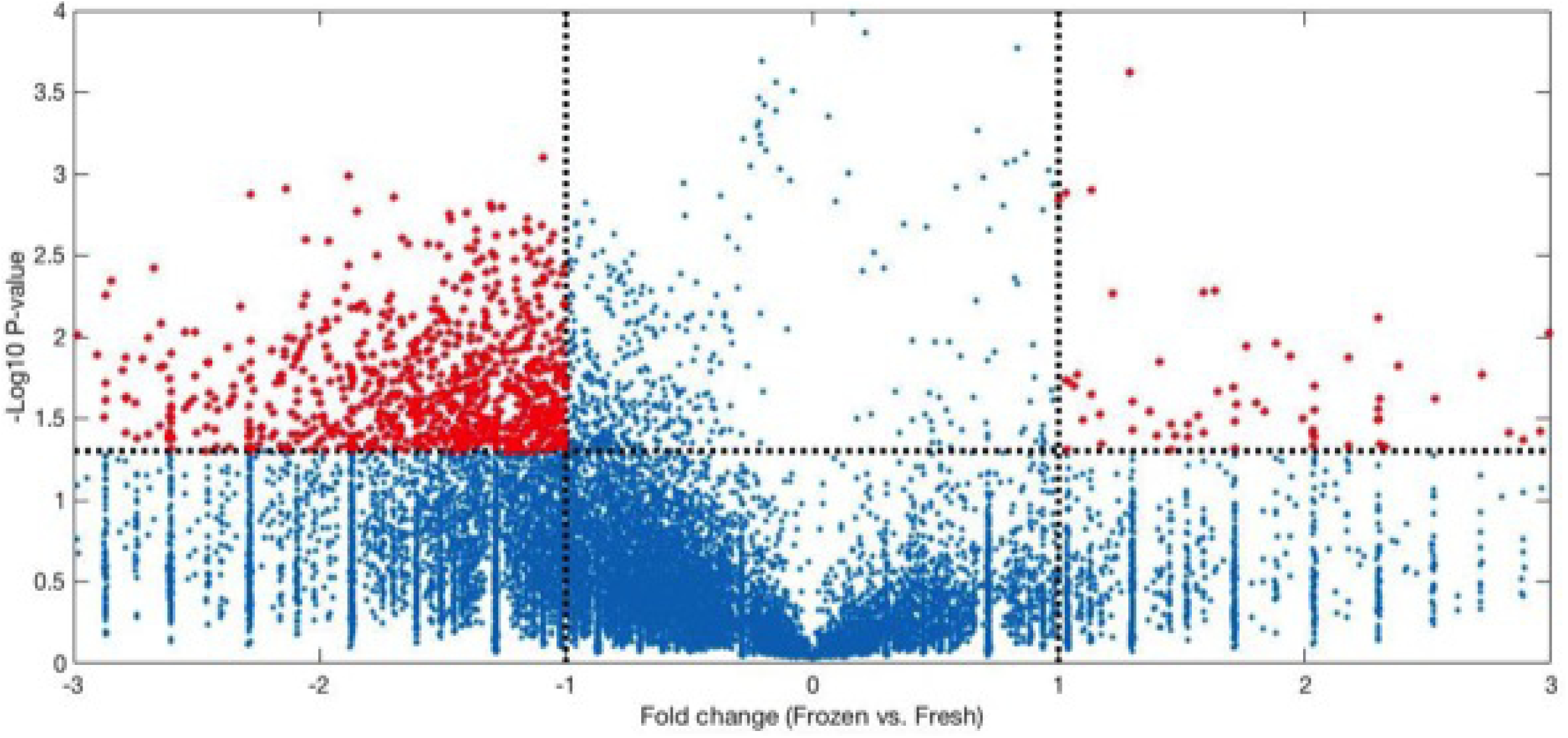
Vulcano plot showing TPM comparison between fresh and frozen tissue at 12 days. Each point represents a gene, characterized in the X-axis as its TPM value in the fresh tissue and in the Y-axis as its TPM value in the frozen tissue. Circled genes represent differentially expressed genes in the two conditions.

Of the 100 transcripts showing increased abundance, 23 could be aligned to the genome build (Supplementary Table 1). These included the *progesterone receptor membrane component (8PGRMC1).* Enriched biological processes (Fig 2b) included extracellular region genes *defensing beta 119, insulin like 3, prostaglandin D2 synthase* and *uteroglobin;* genes associated with negative regulation of cysteine type endopeptidase activity involved in apoptotic processes including *nuclear receptor subfamily 4 group A member* and *paired box 2;* and genes involved in skeletal muscle cell differentiation including *activating transcription factor 3* and *nuclear receptor subfamily 4 group4 A member (Fig 2b)*. STRING analysis revealed no significant enrichments in functional networks for transcripts with increased abundance.

**Figure 2.**
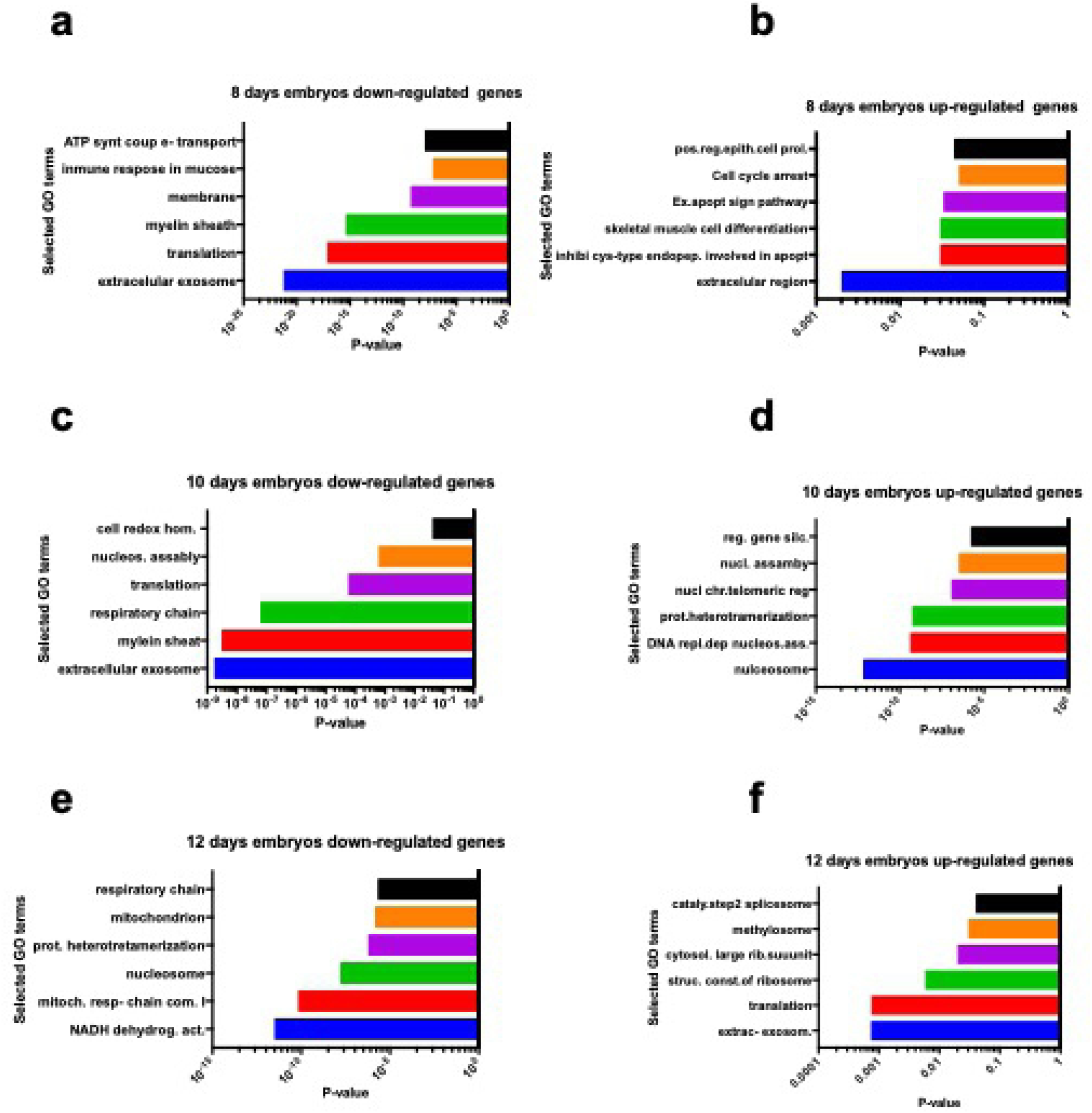
Selected enriched GO terms differentially regulated in equine embryos obtained with fresh and frozen thawed sperm

Transcripts showing decreased abundance provided more information, with 129 transcripts annotated in the equine database. The complete list of transcripts is presented in Supplementary Table 2. Due to the large number of genes retrieved, the threshold was reset at *P* < 0.001 and 62 transcripts were then retrieved (Table 1). Related gene ontology terms are shown in Fig 2a. STRING analysis, performed using a threshold of 0.700, obtained a protein-protein interaction (PPI) enrichment *P* value of < 1.0 x10^-16^ (Fig 4). The complete list of genes in this network with their clustering is presented in Supplementary File 3. Enriched biological processes included *cellular process, iron ion transport, cellular iron ion homeostasis, metabolic process, response to inorganic substance, biological regulation, single-organism process, cellular macromolecule metabolic process, single organism cellular process, cellular metabolic process, response to stimulus, cellular response to zinc ion, transport, regulation of biological process, oxidation-reduction process, cellular component disassembly, cellular nitrogen compound metabolic process, translation, single organism transport, gene expression, positive regulation of nitrogen compound metabolic process, biological process, protein folding, cellular component organization, regulation of cell proliferation, and primary metabolic process*. In addition, enriched terms in KEGG (Kyoto encyclopedia of gene and genomes) pathways included ribosome, Parkinson disease and oxidative phosphorylation (Fig 3a).

**Table 1.** - Enriched biological processes from DEGs (downregulated) in 8 days embryos obtained after AI with frozen thawed sperm, as identified by DAVID functional annotation analysis

| Functional terms of overrepresented biological processes <sup>a</sup> | P value <sup>b</sup> |
| --- | --- |
| Chromosome (21, 35.78) | 7.1 x10<sup>-26</sup> |
| Nucleosome core (19, 42.08) | 9.2 x10<sup>-25</sup> |
| Extracellular exosome (62, 3.56) | 5.7 x10<sup>-22</sup> |
| Histone fold (19, 30.02) | 7.5 x10<sup>-22</sup> |
| Structural constituent of ribosome (24, 13.82) | 5.8 x10<sup>-20</sup> |
| Ribosome (24, 12.89) | 1.21 x10<sup>-19</sup> |
| Histone core (15, 36.91) | 1.69 x10<sup>-18</sup> |
| Translation (21, 14.65) | 7.15 x10<sup>-18</sup> |
| Myelin sheath (18, 16.63) | 3.87 x10<sup>-16</sup> |
| Nucleosome (13, 31.72) | 4.20 x10<sup>-15</sup> |
| Ribonucleoprotein (14, 24.48) | 1.15 x10<sup>-14</sup> |
| Ribosomal protein (13, 29.28) | $1.39 \times 10^{-14}$ |
| Nucleosome assembly (13, 23.18) | $2.14 \times 10^{-13}$ |
| Poly (A) RNA binding (34, 4.29) | $4.69 \times 10^{-13}$ |
| Nuclear nucleosome (10, 40.67) | $1.37 \times 10^{-12}$ |
| Parkinson's disease (18, 9.52) | $2.60 \times 10^{-12}$ |
| Cytosolic small ribosomal subunit (10, 33.98) | $8.65 \times 10^{-12}$ |
| Cytosolic large ribosomal subunit (11, 24.85) | $1.46 \times 10^{-11}$ |
| Systemic lupus erythematosus (16, 10.14) | $2.51 \times 10^{-11}$ |
| Hungtinton's disease (19, 7.38) | $4.19 \times 10^{-11}$ |
| H2B (8, 52.06) | $7.18 \times 10^{-11}$ |
| Nucleus (26, 4.79) | $8.79 \times 10^{-11}$ |
| Oxidative phosphorylation (16, 9.01) | $1.43 \times 10^{-10}$ |
| Histone H2B (8, 48.75) | $1.48 \times 10^{-10}$ |
| DNA binding (21, 6.06) | $1.81 \times 10^{-10}$ |
| Membrane (27, 4.21) | $4.82 \times 10^{-10}$ |
| Alzheimer disease (17, 7.38) | $6.32 \times 10^{-10}$ |
| Alcoholism (16, 7.17) | $3.64 \times 10^{-9}$ |
| ATP synthesis coupled proton transport (7, 41.39) | $1.0 \times 10^{-8}$ |
| Innate immune response in mucose (6, 51.80) | $5.89 \times 10^{-8}$ |
| Antibacterial humoral response (6, 48.15) | $9.1 \times 10^{-8}$ |
| Focal adhesion (15, 6.10) | $1.38 \times 10^{-7}$ |
| DNA binding (18, 4.18) | $1.02 \times 10^{-6}$ |
| DNA replication dependent nucleosome assembly (6, 30.64) | $1.13 \times 10^{-6}$ |
| Hydrogen ion transport (5, 55.38) | $1.38 \times 10^{-6}$ |
| Protein heterotetramerization (6, 29.31) | $1.44 \times 10^{-6}$ |
| Proton transporting ATP synthase activity, rotational mechanism (5, 51.26) | $1.66 \times 10^{-6}$ |
| Cytoplasm (11, 5.91) | $1.60 \times 10^{-5}$ |
| Cytoplasmatic translation (5, 29.56) | $2.00 \times 10^{-5}$ |
| H4 (4, 60.63) | $2.86 \times 10^{-5}$ |
| Viral carcinogenesis (13, 4.39) | $3.09 \times 10^{-5}$ |
| Cardiac muscle contraction (8, 8.53) | $3.57 \times 10^{-5}$ |
| Histone H4 (4, 56.81) | 3.68 x10<sup>-5</sup> |
| Histone H4 conserved site (4, 56.81) | 3.68 x10<sup>-5</sup> |
| TAF (4, 56.88) | 5.56 x10<sup>-5</sup> |
| Defense response to gram positive bacterium (6, 49.69) | 6.14 x10<sup>-5</sup> |
| Tata Box binding protein associated factor (TAF) (4, 14.04) | 7.14 x10<sup>-5</sup> |
| Negative regulation of megakaryocyte differentiation (4, 46.53) | 1.04 x10<sup>-4</sup> |
| ATP hydrolysis coupled ion transport (5, 40.85) | 1.14 x10<sup>-4</sup> |
| Acetylation (6, 19.37) | 1.32 x10<sup>-4</sup> |
| H2A (4, 12.08) | 3.67 x10<sup>-4</sup> |
| Mitochondrial electron transport, cytochrome c to oxygen (3, 27.33) | 4.57 x10<sup>-4</sup> |
| Ribosomal large subunit assembly (4 , 84.27) | 4.94 x10<sup>-4</sup> |
| DNA replication independent nucleosome assembly (4, 24.96) | 4.94 x10<sup>-4</sup> |
| Histone H2A (4, 24.37) | 5.44 x10<sup>-4</sup> |
| V-ATPase proteolipid subunit C-like domain (3, 76.78) | 5.86 x10<sup>-4</sup> |
| DNA templated transcription, initiation (4, 22.47) | 6.81 x10<sup>-4</sup> |
| Nuclear chromosome, telomeric region (6, 8.22) | 7.79 x10<sup>-4</sup> |
| Non alcoholic fatty liver disease (NAFLD) (9, 4.40) | 8.75 x10<sup>-4</sup> |
| Lactate/malate dehydrogenase (3, 63.99) | 8.74 x10<sup>-4</sup> |
| Lactate malate dehydrogenase, N –terminal (3, 63.99) | 8.74 x10<sup>-4</sup> |
| Mitochondrial proton transporting ATP synthase complex (3, 61.00) | 9.60 x10<sup>-4</sup> |

<sup>a</sup> Values in parenthesis represent the number of genes involved in and the fold enrichment of the corresponding functional terms
<sup>b</sup> EASE score examine the significance of gene term enrichment with a modified Fisher's exact test

**Table 2.** Selected enriched Kyoto Encyclopedia of genes and genomes (KEEG) pathways enriched in downregulated transcripts of in 10 days embryos obtained after AI with frozen thawed sperm

| KEEG pathway | Pathway description | observed count | gene false discovery rate |
| --- | --- | --- | --- |
| 190 | Oxidative phosphorylation | 21 | 2,45E-23 |
| 5012 | Parkinson s disease | 20 | 4,64E-21 |
| 5010 | Alzheimer s disease | 15 | 1,05E-12 |
| 5016 | Huntington s disease | 15 | 3,42E-12 |
| 1100 | Metabolic pathways | 28 | 4,29E-09 |
| 3010 | Ribosome | 11 | 8,96E-09 |
| 4260 | Cardiac muscle contraction | 7 | 2,56E-06 |
| 4932 | Non-alcoholic fatty liver disease (NAFLD) | 9 | 3,96E-06 |
| 4141 | Protein processing in endoplasmic reticulum | 9 | 1,61E-05 |

**Figure 3.**
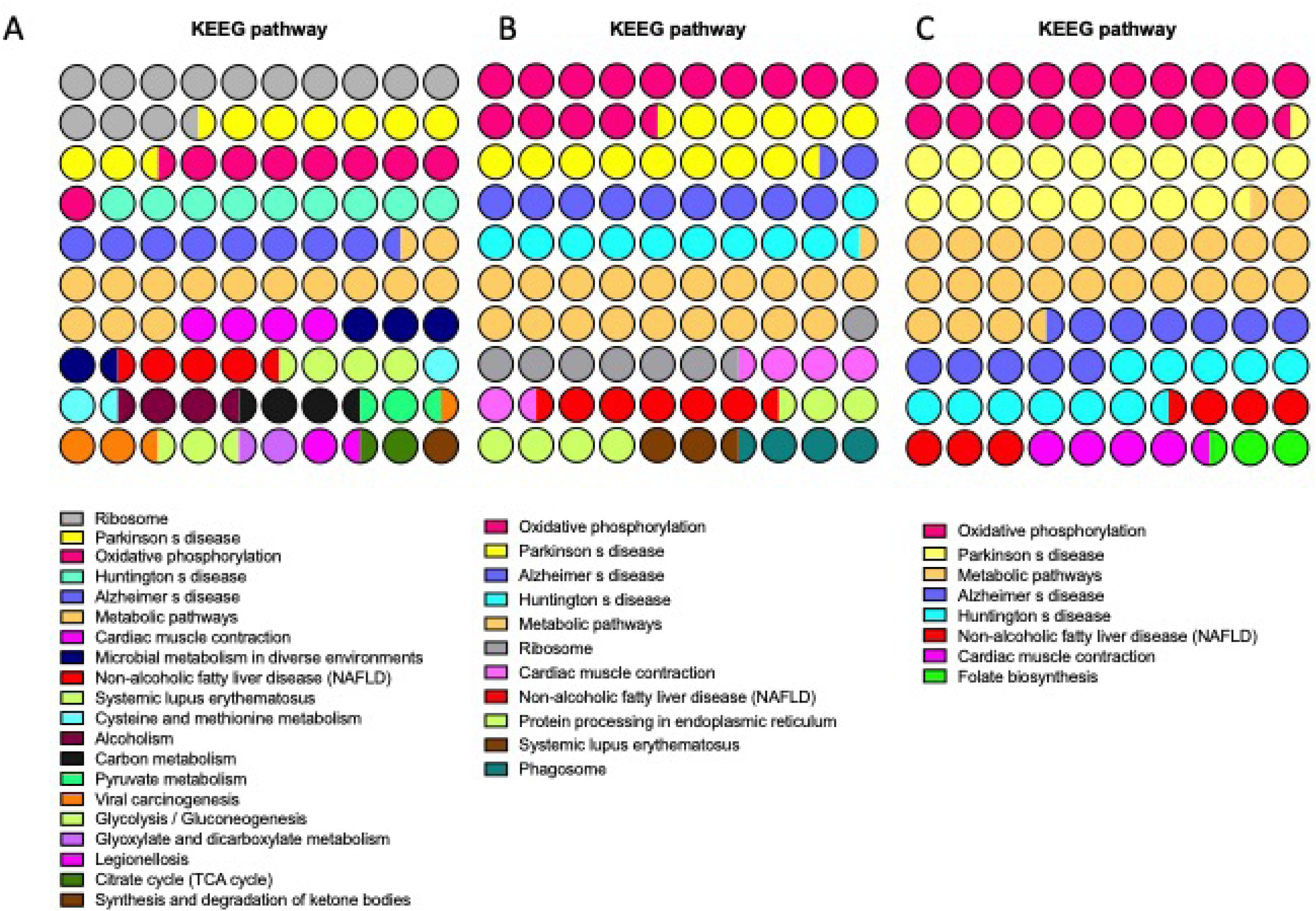
Enriched Kyoto Encyclopedia of genes and genomes (KEEG) pathways in transcripts downregulated in 8 (A) 10 (B) and 12 (C) days embryos obtained with frozen thawed spermatozoa

**Figure 4.**
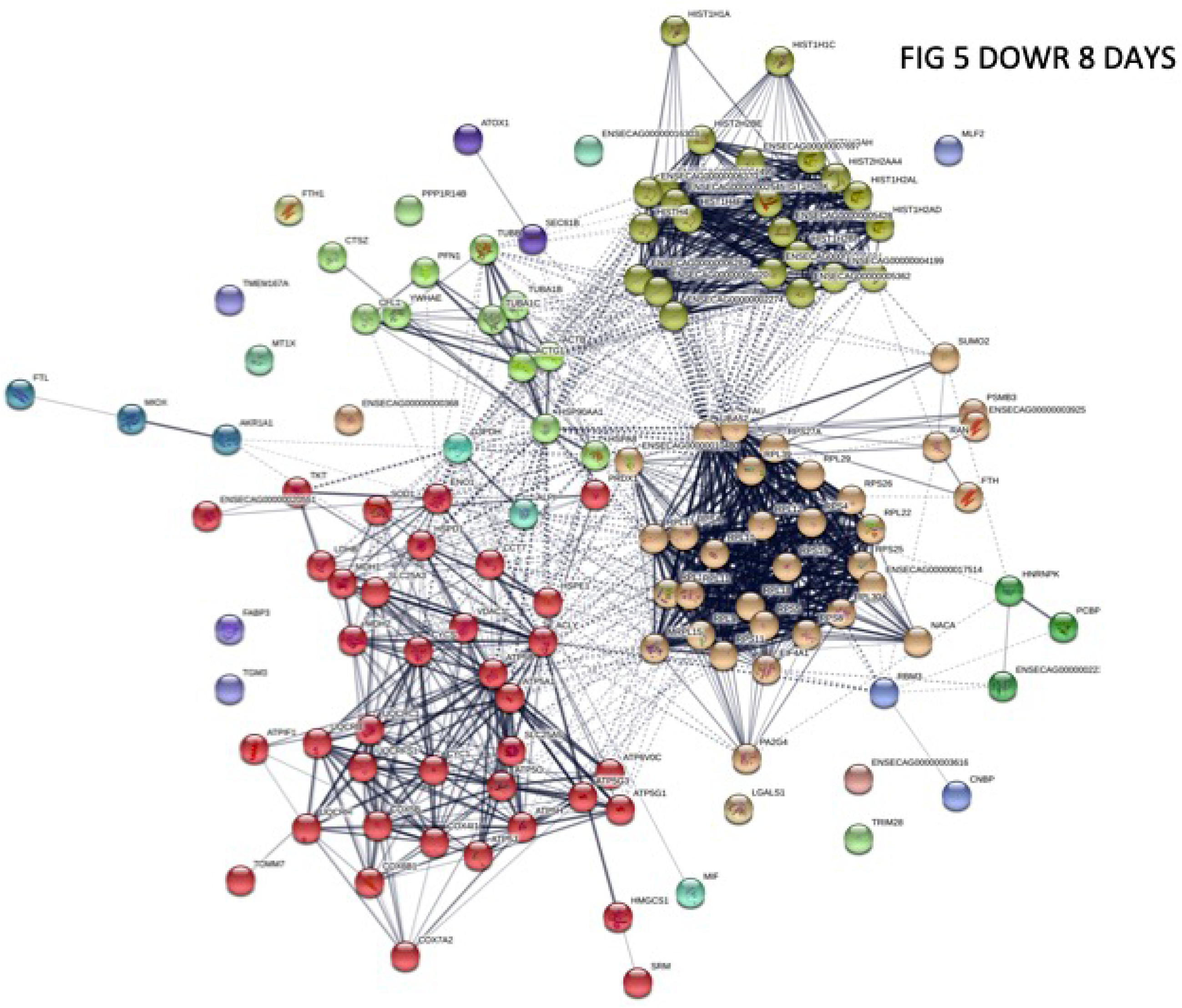
Functional networks (STRING) of transcripts downregulated in 8 days equine embryos obtained with frozen thawed sperm. Functional networks apply to Histones and mitochondrial proteins. Controls are same age embryos from the same mare and stallion obtained with fresh semen

### Day-10 embryos

In Day-10 embryos 239 transcripts showed increased abundance (*P* < 0.01), and 206 showed decreased abundance, in CRYO embryos in comparison with FRSH embryos.

Of the 239 transcripts showing increased abundance, 53 aligned to the genome build (Supplementary Table 4). Functional annotation revealed these genes to be related to the GOterms and KEEG pathways *nucleosome, systemic lupus erythematatosus, DNA replication-dependent nucleosome assembly, protein heterodemerization, alcoholism, nuclear chromosome, telomeric region, regulation of gene silencing, nucleosomal DNA binding, membrane, translation, poly (A) RNA binding, viral carcinogenesis, negative regulation of megakaryocyte differentiation, DNA replication independent nucleosome assembly, extracellular exosome, DNA-templated transcription, xenophagy, ribosome, positive regulation of defense to virus by host, DNA binding, mitochondrion, cytosolic large ribosomal subunit, extracellular space, transcriptional misregulation in cancer, innate immune response in mucosa, U1 snRNP, antibacterial humoral response, telomerase RNA binding* and *mitochondrial small ribosomal subunit* (Fig2 d). STRING analysis revealed a PPI enrichment P value of < 1.0 x10^-16^. Functional enrichment included the PFAM protein domain *Core histone H2A/H2B/H3/HA* and the INTERPRO protein domains, including Histone fold, Histone H3/CNEP-A, Histone H2A/H2B/H3, Histone H4, Histone H4 conserved site, TATA box binding protein associated factor (TAF) and ribosomal protein L23/L15e core domain.

Of the 206 transcripts showing decreased abundance in CRYO embryos at Day 10, 115 were aligned. Enriched KEEG pathways that were also detected in 8-Day embryos (Table 3) included *oxidative phosphorylation, Parkinson disease, Alzheimer disease, Hungtington disease, Metabolic pathways, Ribosome, cardiac muscle contraction*, and *non-alcoholic fatty liver disease.* Three new enriched pathways, *protein processing in endoplasmic reticulum, systemic lupus erythematosus* and *phagosome*, were detected (Fig 3b). More significantly represented GOterms were *ATP synthesis coupled proton transport, translation, nucleosome assembly, cell redox homeostasis, extracellular exosome, myelin sheath, respiratory chain, mitochondrion, extracellular space, NADH dehydrogenase (ubiquinone) activity, structural constituent of ribosome*, and *proton transporting ATP synthase activity rotational mechanism* (Fig 2c). A complete list of enriched GOterms retrieved are given in Table 4. STRING analysis revealed functional networks with a PPI enrichment P value of < 1.0 x10^-16^ (Fig 5). Functional enrichment included the PAFM domains *core histone H2A/H2B/H3/H4, thiorredoxin, NADH deshidrogenase, NADH-Ubiquinone* and *plastoquinone (Complex I), various chains.*

**Table 3.** - Gene ontology annotations enriched in downregulated transcripts of 10 days embryos obtained after AI with frozen thawed sperm

| Term | P Value |
| --- | --- |
| GO:0070062~extracellular exosome | 1,72E-09 |
| GO:0043209~myelin sheath | 3,03E-09 |
| GO:0070469~respiratory chain | 6,30E-08 |
| GO:0022625~cytosolic large ribosomal subunit | 5,33E-07 |
| GO:0008137~NADH dehydrogenase (ubiquinone) activity | 8,07E-07 |
| GO:0005747~mitochondrial respiratory chain complex I | 3,12E-06 |
| GO:0015986~ATP synthesis coupled proton transport | 4,78E-06 |
| GO:0003735~structural constituent of ribosome | 1,66E-05 |
| GO:0000788~nuclear nucleosome | 2,25E-05 |
| GO:0046933~proton-transporting ATP synthase activity, rotational mechanism | 2,90E-05 |
| GO:0005925~focal adhesion | 4,41E-05 |
| GO:0006412~translation | 5,75E-05 |
| GO:0046961~proton-transporting ATPase activity, rotational mechanism | 1,59E-04 |
| GO:0000786~nucleosome | 1,73E-04 |
| GO:0042773~ATP synthesis coupled electron transport | 2,22E-04 |
| GO:0006334~nucleosome assembly | 5,99E-04 |
| GO:0005743~mitochondrial inner membrane | 0,001576512 |
| GO:0004129~cytochrome-c oxidase activity | 0,002995144 |
| GO:0005739~mitochondrion | 0,00444262 |
| GO:0006336~DNA replication-independent nucleosome assembly | 0,005364717 |
| GO:0045261~proton-transporting ATP synthase complex, catalytic core F(1) | 0,011191606 |
| GO:0015991~ATP hydrolysis coupled proton transport | 0,013628447 |
| GO:0006457~protein folding | 0,017943471 |
| GO:0051603~proteolysis involved in cellular protein catabolic process | 0,019505148 |
| GO:0003677~DNA binding | 0,022056606 |
| GO:0006123~mitochondrial electron transport, cytochrome c to oxygen | 0,024396132 |
| GO:0044822~poly(A) RNA binding | 0,032707627 |
| GO:0005753~mitochondrial proton-transporting ATP synthase complex | 0,0332053 |
| GO:0005615~extracellular space | 0,040768468 |
| GO:0005687~U4 snRNP | 0,044030068 |
| GO:0045454~cell redox homeostasis | 0,047994139 |
| GO:0006122~mitochondrial electron transport, ubiquinol to cytochrome c | 0,048204941 |
| GO:1902166~negative regulation of intrinsic apoptotic signaling pathway in response to DNA damage by p53 class mediator | 0,054067016 |
| GO:0006120~mitochondrial electron transport, NADH to ubiquinone | 0,059893472 |
| GO:0045653~negative regulation of megakaryocyte differentiation | 0,065684522 |
| GO:0030330~DNA damage response, signal transduction by p53 class mediator | 0,065684522 |
| GO:0016020~membrane | 0,071097195 |
| GO:0004185~serine-type carboxypeptidase activity | 0,07400431 |
| GO:0034719~SMN-Sm protein complex | 0,075791838 |
| GO:0005685~U1 snRNP | 0,075791838 |
| GO:0002227~innate immune response in mucosa | 0,077161252 |
| GO:0007569~cell aging | 0,077161252 |
| GO:0005682~U5 snRNP | 0,080983251 |
| GO:0071157~negative regulation of cell cycle arrest | 0,082847353 |
| GO:0019731~antibacterial humoral response | 0,082847353 |
| GO:0005975~carbohydrate metabolic process | 0,083234703 |
| GO:0005686~U2 snRNP | 0,086145885 |
| GO:0030970~retrograde protein transport, ER to cytosol | 0,088498889 |
| GO:0000784~nuclear chromosome, telomeric region | 0,089088094 |
| GO:0045787~positive regulation of cell cycle | 0,094116067 |

**Table 4.**
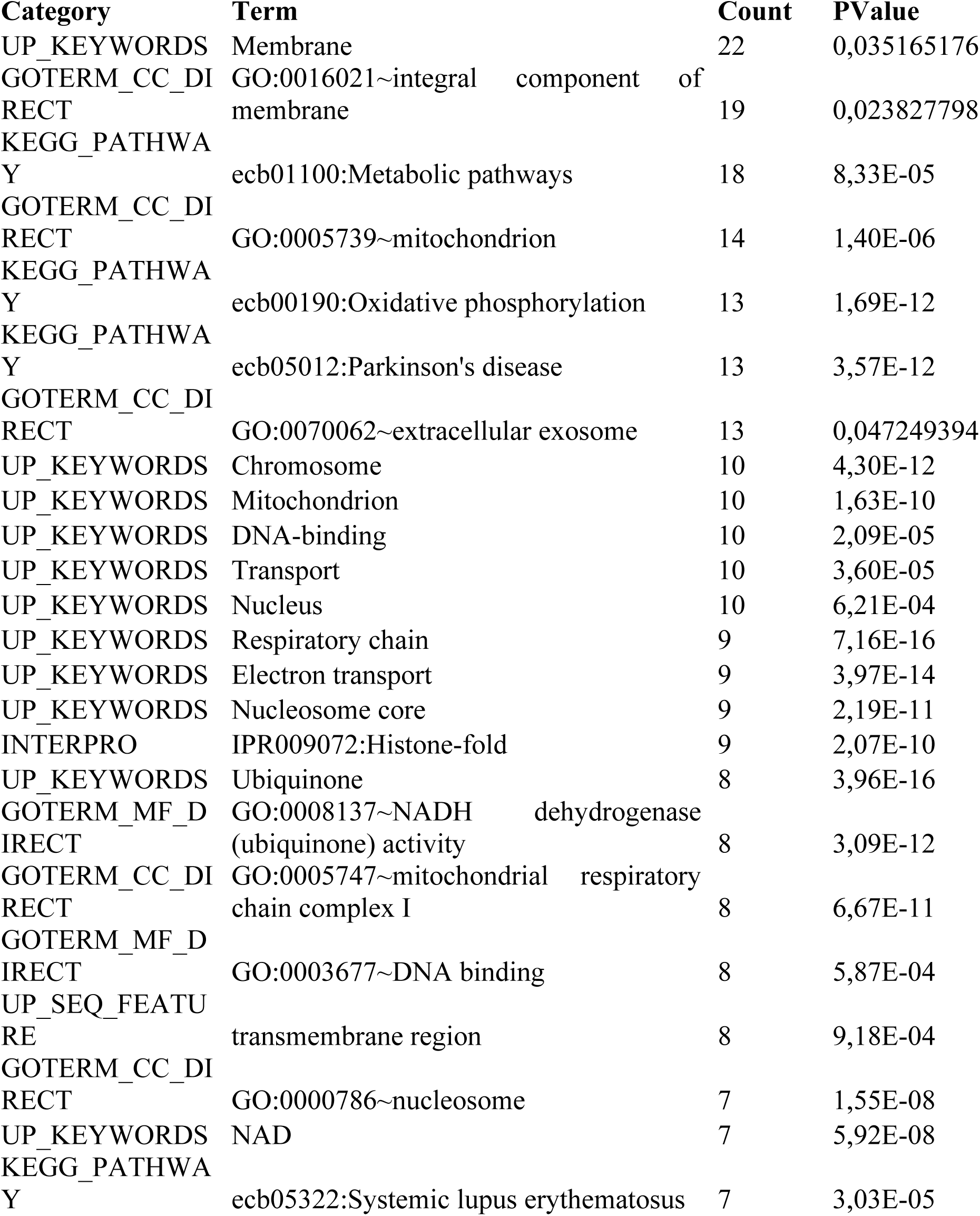
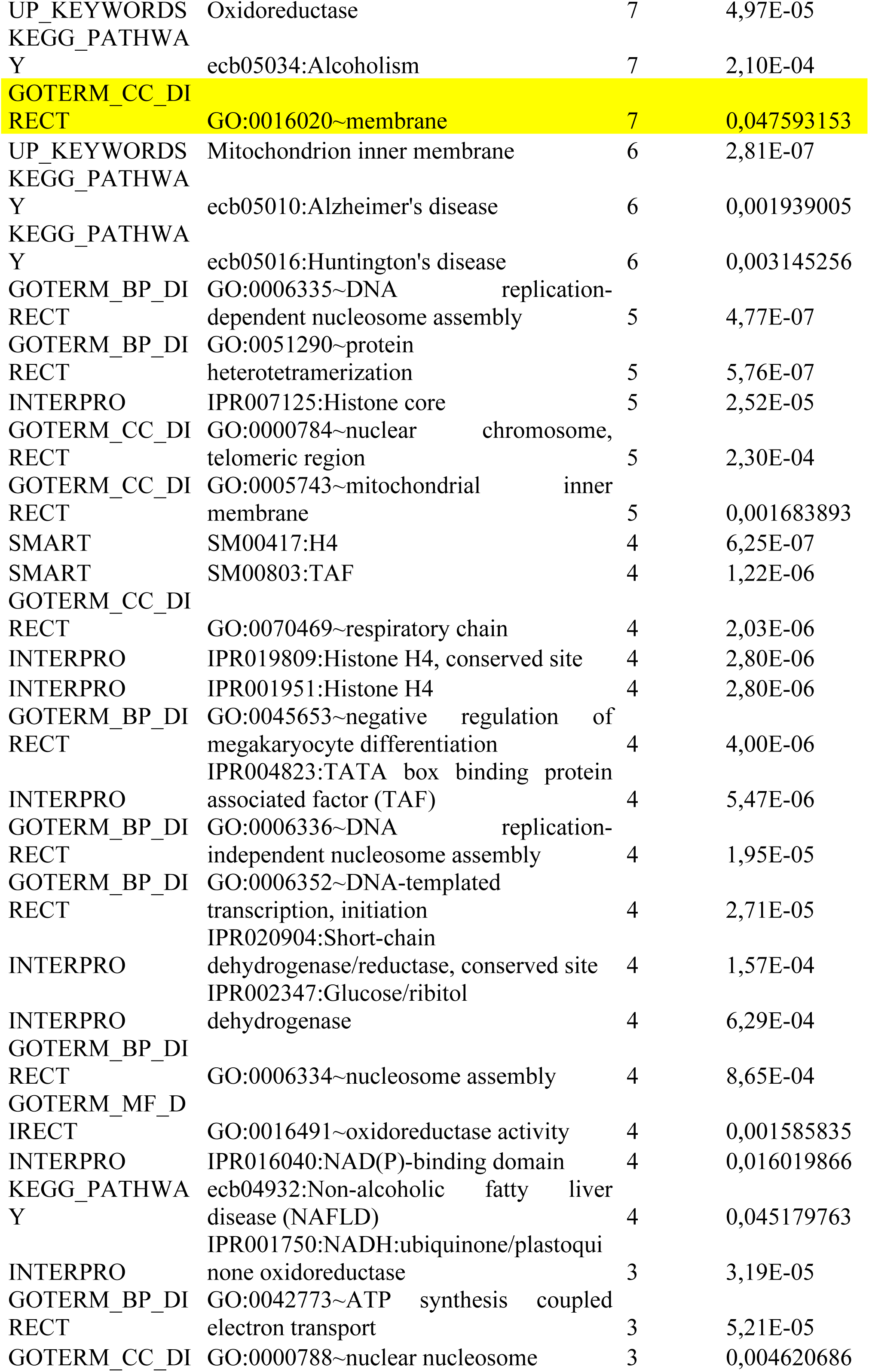
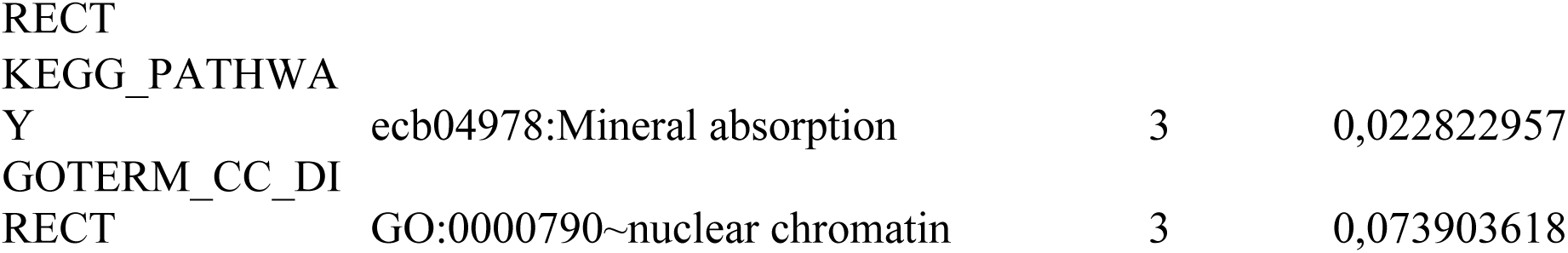
Functional annotation chart of DEGs (downregulated) in 12 days equine embryos obtained after AI with frozen thawed sperm.

**Figure 5.**
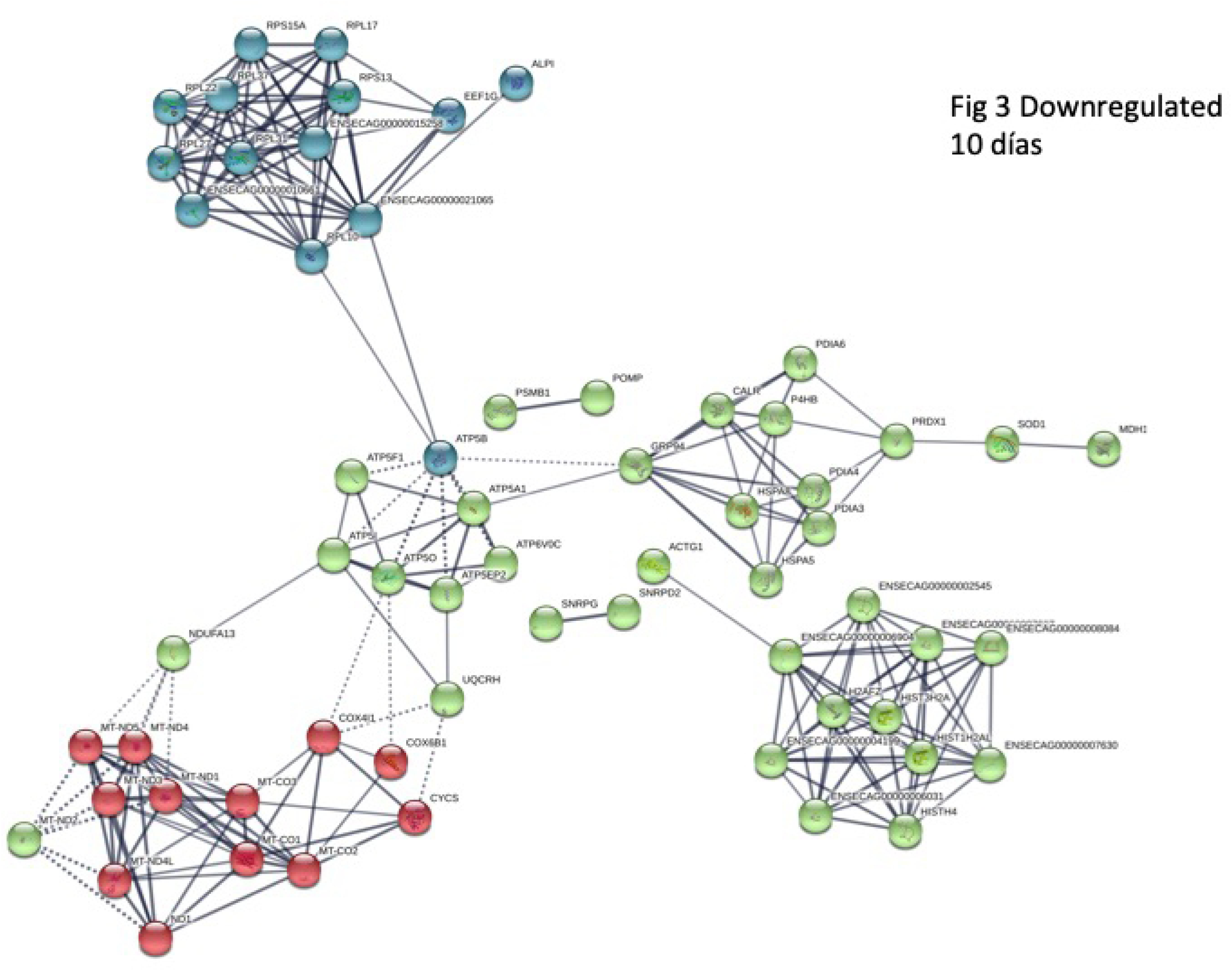
Functional networks (STRING) of transcripts downregulated in 10 days equine embryos obtained with frozen thawed sperm. Functional networks apply to Histones and mitochondrial proteins. Controls are same age embryos from the same mare and stallion obtained with fresh semen

### Day-12 embryos

In Day-12 embryos, 149 transcripts showed increased abundance and 157 showed decreased abundance in CRYO embryos. Of the 149 transcripts with increased abundance, 61 were annotated (Supplementary Table 5). Enriched KKEG pathways included *ribosome* and *Parkinson disease* and the GOterms *extracellular exosome, translation, structural constituent of ribosome, nuclear nucleosome, mitochondrial respiratory chain complex I, cytosolic large ribosomoal subunit, nucleosome assembly, methylosome*, and *catalytic step 2 spliceosome*. On STRING analysis, a PPI enrichment P value of < 8 x10^-10^ was obtained.

Of the 157 transcripts showing decreased abundance in Day-12 CRYO embryos, 60 transcripts aligned to the genome build (Supplementary Table 6). Enriched KEEG pathways, also detected in 8- and 10-day embryos, included *oxidative phosphorylation, Parkinson disease, metabolic pathways, Alzheimer disease, Huntington disease, non-alcoholic fatty acid liver disease* and *cardiac muscle contraction.* In addition a new pathway, *folate biosynthesis*, was enriched (see Fig. 3 for comparative enriched KEEG pathways for transcripts with decreased abundance in 8-, 10- and 12-day CRYO embryos. GOterms enriched annotations (Fig 2e) were *NADH dehydrogenase (ubiquinone) activity, mitochondrial respiratory chain complex I, nucleosome, DNA replication dependent nucleosome assembly, protein heterotetramerization, mitochondrion, respiratory chain, negative regulation of megakaryocyte differentiation, DNA template transcription initiation, ATP synthesis coupled electron transport, nuclear chromosome telomeric region, DNA binding, oxireductase activity, mitochondrial inner membrane, integral component of membrane, mitochondrial electron transport NADH to ubiquinone*, and *extracellular exosome.* The complete list is given in Table 4

STRING analysis revealed functional networks with a PPI enrichment P value of < 1.0 x10^-16^ (Fig 6). Functional enrichment included the PFAM protein domains *core histone H2A/H2B/H3/H4, NADH dehydrogenase*, and *NADH-ubiquinone/plastoquinone (complex I various chains).*

**Figure 6.**
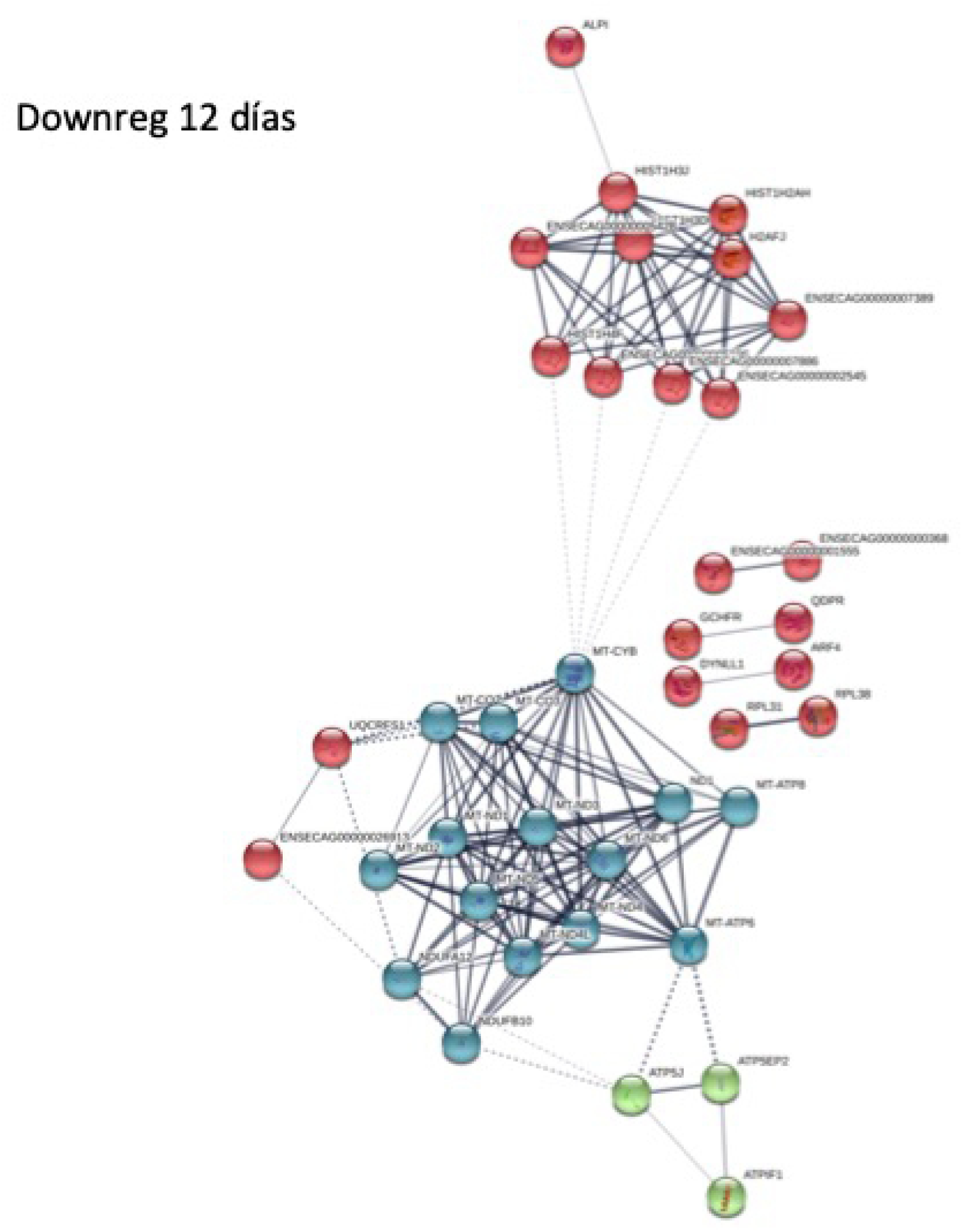
Functional networks (STRING) of transcripts downregulated in 12 days equine embryos obtained with frozen thawed sperm. Functional networks apply to Histones and mitochondrial proteins Controls are same age embryos from the same mare and stallion obtained with fresh semen

### Comparison of downregulated genes with the mouse genome database

In order to explore mechanisms that may relate to reduced viability in embryos obtained using cryopreserved semen, the Mouse Genome Database [27, 28] was queried to determine whether genes downregulated in CRYO equine embryos were orthologs to mouse genes with known associations with embryo lethality.

## Day-8 embryos

In Day-8 CRYO embryos, transcripts of genes associated with the following terms were found to be of low abundance: *failure of zygotic division, decreased embryo size, abnormal embryo size, embryonic growth arrest, embryonic growth retardation, embryonic lethality before implantation-complete penetrance, embryonic lethality between implantation and somite formation-complete penetrance, embryonic lethality between somite formation and embryo turning-complete penetrance, embryonic lethality prior to tooth bud stage, abnormal embryonic tissue morphology, abnormal extraembryonic tissue morphology, delayed allantois development, perinatal lethality incomplete penetrance, prenatal lethality-complete penetrance, preweaning lethality-complete penetrance, abnormal male germ cell apoptosis, abnormal spermatogenesis, azoospermia, male infertility and female infertility.*

### Day 10 embryos

In Day-10 CRYO embryos, the following gene associations cited above for Day-8 embryos were found: *decreased embryo size, abnormal embryo size, failure of zygotic cell division, embryonic lethality between implantation and somite formation, embryonic lethality between implantation and somite formation-complete penetrance, embryonic lethality prior to tooth bud stage, prenatal lethality-complete penetrance, perinatal lethality-incomplete penetrance, preweaning lethality, preweaning lethality-complete penetrance*, and *abnormal spermatogenesis.*

In addition, the following associations were found: *abnormal blastocyst morphology, absent blastocele, abnormal inner cell mass morphology, absent inner cell mass proliferation, empty decidua capsularis, embryonic growth retardation, failure of blastocyst to hatch from the zona pellucida, abnormal preimplantation embryo development, failure to gastrulate, embryonic lethality prior to embryogenesis, failure of embryo implantation, abnormal decidua basalis morphology, abnormal extraembryonic endoderm formation, prenatal lethality prior to heart atrial septation, decreased fetal size, preweaning lethality incomplete penetrance, abnormal gametogenesis, abnormal spermatid morphology, abnormal vas deferens morphology, decreased mature ovarian follicle number, reduced female fertility* and *small ovary.*

### Day 12 embryos

In 12-day CRYO embryos the following gene associations cited above were found: *abnormal embryo size, decreased embryo size, prenatal lethality prior to heart atrial septation, embryonic lethality prior to tooth bud stage, preweaning lethality-complete penetrance, male infertility, female infertility*, and *small ovary.*

In addition, the following associations were found: *incomplete embryo turning, embryonic lethality prior to organogenesis, embryonic lethality during organogenesis-complete penetrance, decreased FSH level, small seminal vesicle, small seminiferous tubules, small testis, absent mature ovarian follicles, abnormal ovulation, abnormal corpus luteum morphology, uterus hypoplasia*, and *vaginal atresia.*

## DISCUSSION

Here we report, for the first time, evidence that procedures performed during handling of sperm, such as freezing and thawing, have a significant impact on critical aspects of the early embryo transcriptome. The equine model used in our study has a number of advantages, including a long pre-attachment embryonic period in which the embryo remains spherical, which facilitates embryo collection, and the possibility of repeated embryo collections from the same animals over successive estrus cycles. Additionally, the stallion serves as an excellent model for the human male, as stallions are typically not selected for sperm quality nor the ability of semen to be cryopreserved, in contrast to males in production species, such as the bull. Moreover, since many stallions reach advanced age, the horse can be used as a model to study the impact of paternal age on embryo quality.

Our study, focused on three embryo ages (8, 10 and 12 days post ovulation), revealed a significant impact of sperm cryopreservation on the transcriptome of the resulting embryo. Importantly, transcripts with decreased abundance reflected genes related to DNA replication and assembly and oxidative phosphorylation. Exploration of differentially-expressed genes at the molecular and cellular level revealed alterations in important functions including ATP synthesis, regulation of transcription, nucleosome assembly, chromatin silencing, protein synthesis, and redox regulation. Alterations in these genes help to explain the reduced fertility observed with cryopreserved sperm attributable to increased early embryo mortality [9, 10].

The pre-implantation period is a period of rapid embryo growth, requiring a ready supply of ATP. The equine embryo appears to have a significant capacity for glycolysis, but also uses oxidative phosphorylation [29]. The KEEG pathways analysis of downregulated genes revealed enriched annotations for oxidative phosphorylation, pyruvate metabolism, glycolysis, and the TCA cycle, suggesting compromised energy metabolism in CRYO embryos. A similar picture was observed in Day-10 and Day-12 embryos, with the pathways for oxidative phosphorylation, metabolic pathways, and non alcoholic fatty liver disease significantly over-represented in transcripts with reduced abundance of all CRYO embryos obtained.

When we evaluated low-abundance equine transcripts for their mouse orthologs, we found that many of the genes downregulated in CRYO embryos have knockout database annotation terms related to reduced embryonic viability. This finding indicates that not only genes related to the metabolism and thus growth of embryos, but also genes directly related to embryo organogenesis, embryo survival, and offspring health are affected by the use of cryopreserved sperm.

While the mechanisms behind the effects reported here are as yet unclear, a major factor may be the well-documented oxidative damage that the genome and epigenome experiences during cryopreservation and thawing [9-12]. Cryopreservation is a major cause of oxidative stress [30] and lipid peroxidation in stallion spermatozoa [8, 14, 31, 32]. Lipid peroxidation in spermatozoa surviving cryopreservation [30] is associated with increased levels of 4-hydroxinonenal (4-HNE) [14]. This compound is able to interact with DNA to form adducts that have been related directly to increased rates of mutation in important cell-cycle regulators [33, 34]. The production of 4-HNE during cryopreservation of stallion spermatozoa is well documented [8, 14, 32], and it is possible that significant amounts of 4-HNE and other toxic lipid aldehydes are incorporated to the oocyte, potentially causing alterations in embryo development. In addition to DNA damage, 4-HNE can alkylate the sperm centrioles, and in horses, as in humans, paternal centrioles are inherited by the embryos. Damaged centrioles may cause disrupted cytoskeletal protein organization during early cleavage [35].

Supporting this line of reasoning, recent reports have linked abnormal early cleavage events and changes in embryo transcript abundance to fertilization with spermatozoa showing oxidative stress. Macaque embryos obtained after fertilization with ROS-treated sperm showed significantly lower rates of development to the four- and eight-cell stages, and changes in transcript abundance for genes related to actin cytoskeleton organization, cell junction assembly and cell adhesion [36]. In our study we also found that genes for cytoskeleton components *tubulin alpha 1 a, tubulin beta 2 class II a* and *actin, cytoplasmic 1, N-terminally processed* were downregulated in 8-day CRYO embryos.

Cryopreservation may also directly affect the epigenome of the paternal DNA; recent studies have shown that cryopreservation increases the level of DNA methylation in equine sperm [10] and the expression of genes important to intracellular regulation of epigenetic status [37]. Notably, we also found significant reduction in abundance of transcripts for histones in CRYO embryos.

The finding that many differentially regulated genes in CRYO embryos are orthologs of mouse genes that have knockout database annotation terms related to reduced embryonic viability provides further evidence linking cryopreserved sperm to reduced embryonic viability. These annotations consistently appeared on analysis of low-abundance transcripts in all CRYO embryos, and included genes related to embryonic growth retardation and embryo lethality. Interestingly, annotations related to male and female infertility were also present; this warrants further investigation on the effect of sperm origin on the fertility of resulting offspring.

In summary, the present study provides for the first time transcriptomic analysis of equine embryos in relation to the handling of semen used for their production. Our data provide strong evidence that cryopreservation of sperm exerts a profound impact on the transcriptome of early embryos. Our findings may stimulate new lines of research to improve this biotechnology in humans and animals

## Acknowledgements

The authors received financial support for this study from the Ministerio de Economía y Competitividad-FEDER, Madrid, Spain, grant <u>AGL2017-83149-R</u>. Junta de Extremadura-FEDER (IB16030 and GR18008) and the The Swedish Research councils VR,(Grant 521-2011-6553) and FORMAS (Grant 2017-00946), Stockholm. JMOR holds a Predoctoral grant from the Valhondo Calaaf Foundation, Cáceres, Spain

## REFERENCES

1. Pena FJ, Garcia BM, Samper JC, Aparicio IM, Tapia JA, Ferrusola CO. Dissecting the molecular damage to stallion spermatozoa: the way to improve current cryopreservation protocols? Theriogenology. 2011;76(7):1177–86. doi: 10.1016/j.theriogenology.2011.06.023. PubMed PMID: 21835453.

2. Dearing CG, Jayasena CN, Lindsay KS. Human sperm cryopreservation in cancer patients: Links with deprivation and mortality. Cryobiology. 2017;79:9– 13. doi: 10.1016/j.cryobiol.2017.10.003. PubMed PMID: 29031884.

3. Jiang XP, Zhou WM, Wang SQ, Wang W, Tang JY, Xu Z, et al. Multivariate model for predicting semen cryopreservation outcomes in a human sperm bank. Asian J Androl. 2017;19(4):404–8. doi: 10.4103/1008-682X.178488. PubMed PMID: 27080478; PubMed Central PMCID: PMCPMC5507083.

4. Lewis N, Morganti, M., Collingwood, F., Grove-White, D.H., Argo, C.M. Utilization of one dose post ovulation breeding with frozen thawed semen at a commercial artificial insemination center: pregnancy rates and post breeding uterine fluid accumulation in comparison with chilled of fresh semen J Equine Vet Sci. 2015;35:882–7.

5. Jodar M, Selvaraju S, Sendler E, Diamond MP, Krawetz SA, Reproductive Medicine N. The presence, role and clinical use of spermatozoal RNAs. Hum Reprod Update. 2013;19(6):604–24. doi: 10.1093/humupd/dmt031. PubMed PMID: 23856356; PubMed Central PMCID: PMCPMC3796946.

6. Sendler E, Johnson GD, Mao S, Goodrich RJ, Diamond MP, Hauser R, et al. Stability, delivery and functions of human sperm RNAs at fertilization. Nucleic Acids Res. 2013;41(7):4104–17. doi: 10.1093/nar/gkt132. PubMed PMID: 23471003; PubMed Central PMCID: PMCPMC3627604.

7. Davila MP, Munoz PM, Bolanos JM, Stout TA, Gadella BM, Tapia JA, et al. Mitochondrial ATP is required for the maintenance of membrane integrity in stallion spermatozoa, whereas motility requires both glycolysis and oxidative phosphorylation. Reproduction. 2016;152(6):683–94. doi: 10.1530/REP-16-0409. PubMed PMID: 27798283.

8. Pena FJ, Plaza Davila M, Ball BA, Squires EL, Martin Munoz P, Ortega Ferrusola C, et al. The Impact of Reproductive Technologies on Stallion Mitochondrial Function. Reprod Domest Anim. 2015;50(4):529–37. doi: 10.1111/rda.12551. PubMed PMID: 26031351.

9. Kopeika J, Thornhill A, Khalaf Y. The effect of cryopreservation on the genome of gametes and embryos: principles of cryobiology and critical appraisal of the evidence. Hum Reprod Update. 2015;21(2):209–27. doi: 10.1093/humupd/dmu063. PubMed PMID: 25519143.

10. Aurich C, Schreiner B, Ille N, Alvarenga M, Scarlet D. Cytosine methylation of sperm DNA in horse semen after cryopreservation. Theriogenology. 2016;86(5):1347–52. doi: 10.1016/j.theriogenology.2016.04.077. PubMed PMID: 27242182.

11. Valcarce DG, Carton-Garcia F, Riesco MF, Herraez MP, Robles V. Analysis of DNA damage after human sperm cryopreservation in genes crucial for fertilization and early embryo development. Andrology. 2013;1(5):723–30. doi: 10.1111/j.2047-2927.2013.00116.x. PubMed PMID: 23970451.

12. Valcarce DG, Carton-Garcia F, Herraez MP, Robles V. Effect of cryopreservation on human sperm messenger RNAs crucial for fertilization and early embryo development. Cryobiology. 2013;67(1):84–90. doi: 10.1016/j.cryobiol.2013.05.007. PubMed PMID: 23727067.

13. Teperek M, Simeone A, Gaggioli V, Miyamoto K, Allen GE, Erkek S, et al. Sperm is epigenetically programmed to regulate gene transcription in embryos. Genome Res. 2016;26(8):1034–46. doi: 10.1101/gr.201541.115. PubMed PMID: 27034506; PubMed Central PMCID: PMCPMC4971762.

14. Martin Munoz P, Ortega Ferrusola C, Vizuete G, Plaza Davila M, Rodriguez Martinez H, Pena FJ. Depletion of Intracellular Thiols and Increased Production of 4-Hydroxynonenal that Occur During Cryopreservation of Stallion Spermatozoa Lead to Caspase Activation, Loss of Motility, and Cell Death. Biol Reprod. 2015;93(6):143. doi: 10.1095/biolreprod.115.132878. PubMed PMID: 26536905.

15. Plaza Davila M, Martin Munoz P, Tapia JA, Ortega Ferrusola C, Balao da Silva CC, Pena FJ. Inhibition of Mitochondrial Complex I Leads to Decreased Motility and Membrane Integrity Related to Increased Hydrogen Peroxide and Reduced ATP Production, while the Inhibition of Glycolysis Has Less Impact on Sperm Motility. PLoS One. 2015;10(9):e0138777. doi: 10.1371/journal.pone.0138777. PubMed PMID: 26407142; PubMed Central PMCID: PMCPMC4583303.

16. Timme-Laragy AR, Hahn ME, Hansen JM, Rastogi A, Roy MA. Redox stress and signaling during vertebrate embryonic development: Regulation and responses. Semin Cell Dev Biol. 2017. doi: 10.1016/j.semcdb.2017.09.019. PubMed PMID: 28927759; PubMed Central PMCID: PMCPMC5650060.

17. Fernandez-Gonzalez R, Moreira PN, Perez-Crespo M, Sanchez-Martin M, Ramirez MA, Pericuesta E, et al. Long-term effects of mouse intracytoplasmic sperm injection with DNA-fragmented sperm on health and behavior of adult offspring. Biol Reprod. 2008;78(4):761–72. doi: 10.1095/biolreprod.107.065623. PubMed PMID: 18199884.

18. Bouckenheimer J, Assou S, Riquier S, Hou C, Philippe N, Sansac C, et al. Long non-coding RNAs in human early embryonic development and their potential in ART. Hum Reprod Update. 2016;23(1):19–40. doi: 10.1093/humupd/dmw035. PubMed PMID: 27655590.

19. Chen Q, Yan W, Duan E. Epigenetic inheritance of acquired traits through sperm RNAs and sperm RNA modifications. Nat Rev Genet. 2016;17(12):733–43. doi: 10.1038/nrg.2016.106. PubMed PMID: 27694809; PubMed Central PMCID: PMCPMC5441558.

20. Saunders CM, Larman MG, Parrington J, Cox LJ, Royse J, Blayney LM, et al. PLC zeta: a sperm-specific trigger of Ca(2+) oscillations in eggs and embryo development. Development. 2002;129(15):3533-44. PubMed PMID: 12117804.

21. Fujimoto S, Yoshida N, Fukui T, Amanai M, Isobe T, Itagaki C, et al. Mammalian phospholipase Czeta induces oocyte activation from the sperm perinuclear matrix. Dev Biol. 2004;274(2):370–83. doi: 10.1016/j.ydbio.2004.07.025. PubMed PMID: 15385165.

22. Banrezes B, Sainte-Beuve T, Canon E, Schultz RM, Cancela J, Ozil JP. Adult body weight is programmed by a redox-regulated and energy-dependent process during the pronuclear stage in mouse. PLoS One. 2011;6(12):e29388. doi: 10.1371/journal.pone.0029388. PubMed PMID: 22216268; PubMed Central PMCID: PMCPMC3247262.

23. Iqbal K, Chitwood JL, Meyers-Brown GA, Roser JF, Ross PJ. RNA-seq transcriptome profiling of equine inner cell mass and trophectoderm. Biol Reprod. 2014;90(3):61. doi: 10.1095/biolreprod.113.113928. PubMed PMID: 24478389; PubMed Central PMCID: PMCPMC4435230.

24. Cuervo-Arango J, Aguilar J, Newcombe JR. Effect of type of semen, time of insemination relative to ovulation and embryo transfer on early equine embryonic vesicle growth as determined by ultrasound. Theriogenology. 2009;71(8):1267–75. doi: 10.1016/j.theriogenology.2008.12.020. PubMed PMID: 19246082.

25. Jiao X, Sherman BT, Huang da W, Stephens R, Baseler MW, Lane HC, et al. DAVID-WS: a stateful web service to facilitate gene/protein list analysis. Bioinformatics. 2012;28(13):1805–6. doi: 10.1093/bioinformatics/bts251. PubMed PMID: 22543366; PubMed Central PMCID: PMCPMC3381967.

26. Szklarczyk D, Franceschini A, Wyder S, Forslund K, Heller D, Huerta-Cepas J, et al. STRING v10: protein-protein interaction networks, integrated over the tree of life. Nucleic Acids Res. 2015;43(Database issue):D447–52. doi: 10.1093/nar/gku1003. PubMed PMID: 25352553; PubMed Central PMCID: PMCPMC4383874.

27. Blake JA, Bult CJ, Eppig JT, Kadin JA, Richardson JE, Mouse Genome Database G. The Mouse Genome Database genotypes::phenotypes. Nucleic Acids Res. 2009;37(Database issue):D712–9. doi: 10.1093/nar/gkn886. PubMed PMID: 18981050; PubMed Central PMCID: PMCPMC2686566.

28. Bult CJ, Eppig JT, Kadin JA, Richardson JE, Blake JA, Mouse Genome Database G. The Mouse Genome Database (MGD): mouse biology and model systems. Nucleic Acids Res. 2008;36(Database issue):D724–8. doi: 10.1093/nar/gkm961. PubMed PMID: 18158299; PubMed Central PMCID: PMCPMC2238849.

29. Lane M, O’Donovan MK, Squires EL, Seidel GE, Jr., Gardner DK. Assessment of metabolism of equine morulae and blastocysts. Mol Reprod Dev. 2001;59(1):33–7. doi: 10.1002/mrd.1004. PubMed PMID: 11335944.

30. Ortega Ferrusola C, Gonzalez Fernandez L, Morrell JM, Salazar Sandoval C, Macias Garcia B, Rodriguez-Martinez H, et al. Lipid peroxidation, assessed with BODIPY-C11, increases after cryopreservation of stallion spermatozoa, is stallion-dependent and is related to apoptotic-like changes. Reproduction. 2009;138(1):55–63. doi: 10.1530/REP-08-0484. PubMed PMID: 19380427.

31. Ortega-Ferrusola C, Anel-Lopez L, Martin-Munoz P, Ortiz-Rodriguez JM, Gil MC, Alvarez M, et al. Computational flow cytometry reveals that cryopreservation induces spermptosis but subpopulations of spermatozoa may experience capacitation-like changes. Reproduction. 2017;153(3):293–304. doi: 10.1530/REP-16-0539. PubMed PMID: 27965398.

32. Munoz PM, Ferrusola CO, Lopez LA, Del Petre C, Garcia MA, de Paz Cabello P, et al. Caspase 3 Activity and Lipoperoxidative Status in Raw Semen Predict the Outcome of Cryopreservation of Stallion Spermatozoa. Biol Reprod. 2016;95(3):53. doi: 10.1095/biolreprod.116.139444. PubMed PMID: 27417910.

33. Feng Z, Hu W, Amin S, Tang MS. Mutational spectrum and genotoxicity of the major lipid peroxidation product, trans-4-hydroxy-2-nonenal, induced DNA adducts in nucleotide excision repair-proficient and -deficient human cells. Biochemistry. 2003;42(25):7848–54. doi: 10.1021/bi034431g. PubMed PMID: 12820894.

34. Wei X, Yin H. Covalent modification of DNA by alpha, beta-unsaturated aldehydes derived from lipid peroxidation: Recent progress and challenges. Free Radic Res. 2015;49(7):905–17. doi: 10.3109/10715762.2015.1040009. PubMed PMID: 25968945.

35. Lu Y, Lin M, Aitken RJ. Exposure of spermatozoa to dibutyl phthalate induces abnormal embryonic development in a marine invertebrate Galeolaria caespitosa (Polychaeta: Serpulidae). Aquat Toxicol. 2017;191:189–200. doi: 10.1016/j.aquatox.2017.08.008. PubMed PMID: 28843738.

36. Burruel V, Klooster KL, Chitwood J, Ross PJ, Meyers SA. Oxidative damage to rhesus macaque spermatozoa results in mitotic arrest and transcript abundance changes in early embryos. Biol Reprod. 2013;89(3):72. doi: 10.1095/biolreprod.113.110981. PubMed PMID: 23904511; PubMed Central PMCID: PMCPMC4094196.

37. Zeng C, Peng W, Ding L, He L, Zhang Y, Fang D, et al. A preliminary study on epigenetic changes during boar spermatozoa cryopreservation. Cryobiology. 2014;69(1):119–27. doi: 10.1016/j.cryobiol.2014.06.003. PubMed PMID: 24974820.

